# 4-Methyllumifrone (4-MU) can improve learning and memory after cerebral ischemia/reperfusion injury in rats

**DOI:** 10.1101/2024.11.29.626098

**Authors:** Shaghayegh Tamouk, Hamzeh Mirshekari Jahangiri, Elham Kashafi Jahromi, Michael R Hamblin, Fatemeh Ramezani, Nahid Aboutaleb

## Abstract

**Background:** Stroke is the sixth leading cause of death and lifelong disability for millions of people in the United States. Cerebral ischemia leads to oxidative stress, excitotoxicity, inflammation, apoptosis; additionally, impairments in memory and learning occur in the majority of subjects with ischemic stroke. The lack of definitive treatment has sparked extensive research into novel therapeutic strategies, including the use of 4-methylumbelliferone (4-MU), a coumarin derivative with potential neuroprotective properties. The present study examines the impact of 4-MU on reducing cerebral ischemia-reperfusion (I/R) injury and learning and memory impairments in male Wistar rats.

**Method:** The animals were exposed to middle cerebral artery occlusion (MCAO) and were treated with one dose of 4-MU (at a dosage of 25 mg/kg) dissolved in DMSO 0.9%. Automated shuttle box and Morris water maze (MWM) test were employed to evaluate learning and memory impairments. Western blot assay, TTC staining and Nissl staining were used to measure protein expression, infarct volume, and cell death, respectively.

**Results:** Results showed that treatment with 4-MU reduced infarct volume and improved learning and memory impairments by down-regulating HAS1 and HAS2. 4-MU has been shown to modulate the release of pro-inflammatory cytokines including TNF-α and IL-1β, as well as anti-inflammatory markers like IL-10 and also reduced oxidative stress markers in the brain.

**Conclusion:** Generally, the neuroprotective effects of 4-MU against cerebral I/R injury can be attributed to the down-regulation of HAS1 and HAS2.

## 1. Introduction

One of the main outcomes following ischemic stroke is impairments in memory and learning, which strongly influences patients’ quality of life (1). Previous investigations have shown that cerebral ischemia for a short time (less than 10 minutes) contributes to the initiation of cell death in the hippocampus and causes deficits in learning and memory functions (2).

A large number of individuals who survive from the acute phase of ischemic stroke encounter some adverse effects related to deficits in memory and learning because available treatments are not completely effective (3). According to the clinical studies, immediate or delayed dysfunctions in memory and learning occur in 25–30% of post-stroke survivors which heavily influences rehabilitation programs for performing daily living activities (4, 5). Hyaluronan (HA) is a key ingredient of the brain extracellular matrix and plays a crucial role in processes like angiogenesis, cellular proliferation, differentiation, and migration. Increased synthesis of total HA and low molecular mass 3–10 disaccharides of HA (o-HA) in the serum of patients with ischemic stroke (1, 3, 7 and 14 days) and post-mortem tissues have been reported in a prior investigation, mainly attributed to overexpression of HA synthases (HAS1 and HAS2) and hyaluronidases in inflammatory cells in peri-infarcted areas of the cerebral tissue (6). Elevated expression of hyaluronidase 1 and HAS2 in astrocytes following ischemic stroke aggravates infarct volume and suppressing hyaluronidase activity early after stroke can contribute to functional recovery (7).

4-methylumbelliferone (4-MU), an inhibitor of hyaluronic acid synthesis, has been found to have a protective impact against ischemia/reperfusion injury and other disorders by inhibition of oxidative stress and inflammation (8, 9, 10, 11, 12). The protective impact of 4-MU is associated with reducing macrophage invasion (13).

The accumulating evidence suggests that 4-MU may be a valuable adjunctive therapy for treating ischemic stroke and other serious diseases (14, 15, 16). Its multifaceted mechanism of action, encompassing anti-inflammatory, antioxidant, and anti-apoptotic properties, may provide a therapeutic advantage over single-target agents (17). Ultimately, the development of 4-MU as a therapeutic agent for ischemic stroke may contribute to improved patient outcomes and a reduction in the socioeconomic burden of this devastating disease. Here we tested whether 4-MU can reduce cerebral I/R injury by modulating the activities of HAS1 and HAS2 in a rat model.

## 2. Materials and Methods

### 2.1. Animals and Experimental Design

Wistar rats (male, 275 ± 15 g) were purchased from the Animal Center of Iran University of Medical Sciences (IUMS). Controlled conditions including 12h light-dark schedule at a temperature of 22 ± 2°C, with unrestricted access to standard food and water ad libitum, were selected for keeping the animals. All of the rats underwent 3 days of handling. All protocols and procedures relevant to the animals were authorized by the Ethics Committee of IUMS and were performed under the National Institutes of Health Guide for the Care and Use of Laboratory Animals.

116 rats were randomly assigned into four groups (n=29, per group): Sham, I/R, vehicle, and 4- MU (25 mg/kg). In sham group, the animals were only subjected to surgical shock. In the I/R group, the animals were subjected to MCAO for 30 min and then reperfusion was allowed for 24h. In the vehicle group, the animals were subjected to MCAO for 30 min and at the onset of reperfusion, solvent (DMSO 0.9%) was administered intraperitoneally. In the 4-MU group, the animals were subjected to MCAO for 30 min and at the onset of reperfusion, 4-MU (at a dosage of 25 mg/kg) dissolved in DMSO 0.9% was intraperitoneally administered.

### 2.2. Ischemic Stroke Model (MCAO)

The MCAO/R model was used to induce ischemia as described by Longa et al (18). The animals were anesthetized using a combination of ketamine (80□mg/kg) and xylazine (10□mg/kg) intraperitoneally. After isolation of the right common artery, external carotid artery (ECA), and internal carotid artery (ICA), the distal end of the ECA was cut and a 4–0 silicon rubber-coated monofilament containing a tapered end was inserted into the right ICA and forwarded approximately 18 ± 2 mm to feel slight resistance as an indicator of blocking the MCA. To permit reperfusion, the silicon rubber-coated monofilament was taken out after 30 minutes. Similar surgical procedures were performed for sham animals except inserting the silicon rubber-coated monofilament.

### 2.3. Automated Shuttle Box Test

Seven days after surgery, the rats underwent a shuttle box test. A shuttle box including a two-part plexiglass box with a bright section and a dark section, and the equal dimensions of the two sections (20×20×40 cm) was employed to assess learning and memory function. The instrument includes stainless steel bars on the floor of both sections with a 1 cm distance, a 100-W light bulb 40 cm above the device’s bright side, and a guillotine door between the two sections. The experiment was carried out in 3 steps: 1) Adaption: each rat was habituated with the apparatus for ≥5 min on two consecutive days prior to start of the test. 2) Acquisition: in the third day, each rat was placed in the bright section, and the chamber is kept dark for 2 min. both dark and bright sections are separated by a guillotine door. At the end of the period and after turning on the chamber’s light and opening the guillotine door, the time it takes for the animal to enter from the light to the dark chamber is recorded by a chronometer and described as “Initial latency”. Then, after closing the door, a single electric shock (1 mA for 1 second) is exerted on the rat. One min after the end of the experiment, the rat is transferred back to its cage. At this step, animals with a delay more than 60 seconds are excluded from the experiment. The retention and recall step were done 24 h after the second step on the fourth day. This step is like the prior step, but no shock is given when the rat enters the dark section. STL (step-through latency) is recorded at this point; STL is an indicator of how long the animal stays in the light section before entering the dark section. The cut-off time was 480s if the animal did not move to the dark section.

### 2.4. Morris water maze (MWM) test

Seven days after surgery, the rats underwent a Morris water maze (MWM) test. Spatial learning and memory performance were investigated using the MWM. The maze employed in this study consists of a black circular pool (diameter = 150□cm, height = 60□cm) filled with water at 20□±□1°C to a height of 25□cm. The four quadrants of the black circular pool are arbitrarily described as southeast, and southwest northeast, and northwest. In the target quadrant (southeast), a hidden circular platform (diameter = 10□cm and located 1.5□cm below the water surface) was placed. During all days of the test, visual cues were placed around the maze and recording the swimming paths of rats during trials was performed using a camera fixed above the center of the maze. Before triggering each daily test, rats were placed in a dark test room for 30□min. Four trials each day for three consecutive days were performed for each animal. Each animal was subjected to a 20□s habituation time on the platform just on the first day. The rats were placed in water facing the maze wall and allowed to swim for 90□s until they discovered the platform at the initiation of all trials. The rats that lacked the ability to explore the platform were manually guided to it. After reaching the platform, they were permitted to remain for 20□s on the platform and then returned to their cage. A random selection for triggering positions was done each day. The probe (retention test) and visible test were performed on day 4. To carry out the probe test, the escape platform was removed from the maze; afterward, rats were placed in water from a point opposite to the target quadrant and allowed to swim for up to 60 seconds. Sensorimotor abilities of animals were measured using a visible platform test after the probe trial. Processing and analysis of the recorded behaviors of each rat were done using a computerized system (Noldus EthoVision, version 11.1).

### 2.5. TTC staining

The brain infarct volume was measured using TTC staining. After sacrificing the animals at the end of the experiment, brain tissues were separated and cut into 2 mm coronal segments. To perform staining, the brain sections were immersed in 2% TTC for 15 min at 37 □ and fixed with 4% paraformaldehyde to increase contrast before taking photographs. Calculation of the percentage of infarct volume was done based on the following formula:

Brain infarct volume (%) = [left hemisphere volume − (right hemisphere volume − infarct volume)] / left hemisphere volume × 100 %.

### 2.6. Western blotting

Rat brain tissue proteins were extracted from infarcted regions using the RIPA lysis buffer. A BCA protein assay kit was employed to measure the protein concentrations. After separation of 50 μg protein of each sample based on molecular weight by electrophoresis, they were transferred to PVDF membranes. After placing the membranes in primary antibodies against HAS1 and HAS2 overnight at 4 °C, they were incubated with corresponding HRP-conjugated secondary antibodies for 2 hours at 37 □. The color development of the PVDF membranes was done using the chemiluminescent HRP Substrate (Millipore). Quantification of the protein values was performed using Image-J software. β-actin was employed as the loading control.

### 2.7. Histopathological examinations

Histopathological alterations of cerebral tissues were determined using Nissl staining. The brain tissues were fixed with 4% paraformaldehyde, embedded in paraffin sections, and cut into 5 µm slides before being Nissl stained and sealed. An optical microscope (Motic China Group Co., Ltd., BA210Digital, Xiamen, China) was used to investigate the pathological alterations in the CA1 region of the hippocampus.

### 2.8. Statistical analysis

The findings were defined as mean ±□Standard error of the mean (SEM). Analysis of all data was done by using one-way ANOVA followed by Bonferroni’s test as a post hoc analytical test (GraphPad Prism 9.0 software). P-value less than 0.05 indicates a statistically significant difference.

## 3. Results

### 3.1. Infarct Volume

The obvious cerebral infarction was evident in the I/R cohort compared to the sham, indicating the successful establishment of the stroke model. As expected, a significant reduction in infarct size was observed in MCAO rats treated with 4-MU (25 mg/kg) (Figure 1).

**Figure 1.**
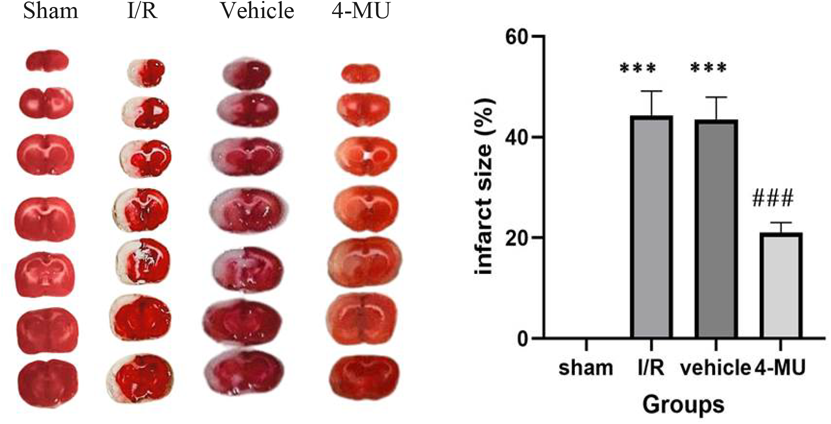
Representative images of TTC staining in different groups. Red color shows healthy regions, while white color demonstrate infarcted regions (n=5 per group). ***p□<□0.001 vs. sham; ### p□<□0.001 vs. I/R and vehicle groups). Data were expressed as mean ±□SEM.

### 3.2. Learning and memory

Remarkable reductions in STL were observed in animals exposed to MCAO compared to the sham. 4-MU (25 mg/kg) could contribute to restoration of STL compared to the I/R and vehicle groups. Moreover, MCAO rats exhibited significantly more time spent in dark chamber than the sham. Treatment with only DMSO 0.9% did not change the time spent in dark chamber, whereas treatment with 4-MU (25 mg/kg) markedly decreased time spent in dark chamber compared to the I/R and vehicle cohorts (Figure 2 A-B).

**Figure 2.**
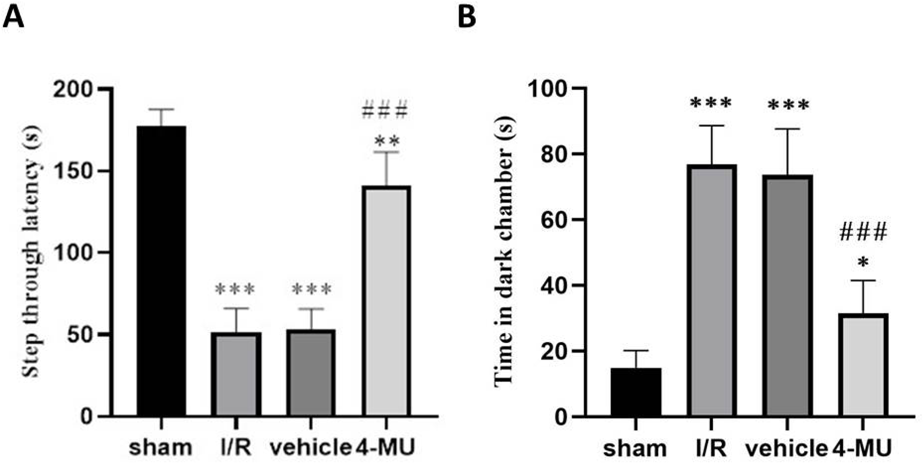
Impact of 4-MU on ischemic stroke-induced learning and memory impairments. **A)** STL, **B)** time in dark chamber (n=7). ***p□<□0.001, **p□<□0.01, and *p□<□0.05 vs. sham; ### p□<□0.001 vs. I/R and vehicle groups). Data were expressed as mean ±□SEM.

### 3.3. Behavioral parameters in the MWM test

Reduced levels of the time spent in the goal quarter and increased latency time were observed in the I/R and vehicle cohorts compared to the sham (Figure 3 A-B). Treatment of I/R rats with 4- MU (25 mg/kg) significantly improved the time spent in the goal quarter and decreased latency time compared to the I/R and vehicle cohorts. There were no significant differences among the various cohorts regarding the speed index (Figure 3 C).

**Figure 3.**
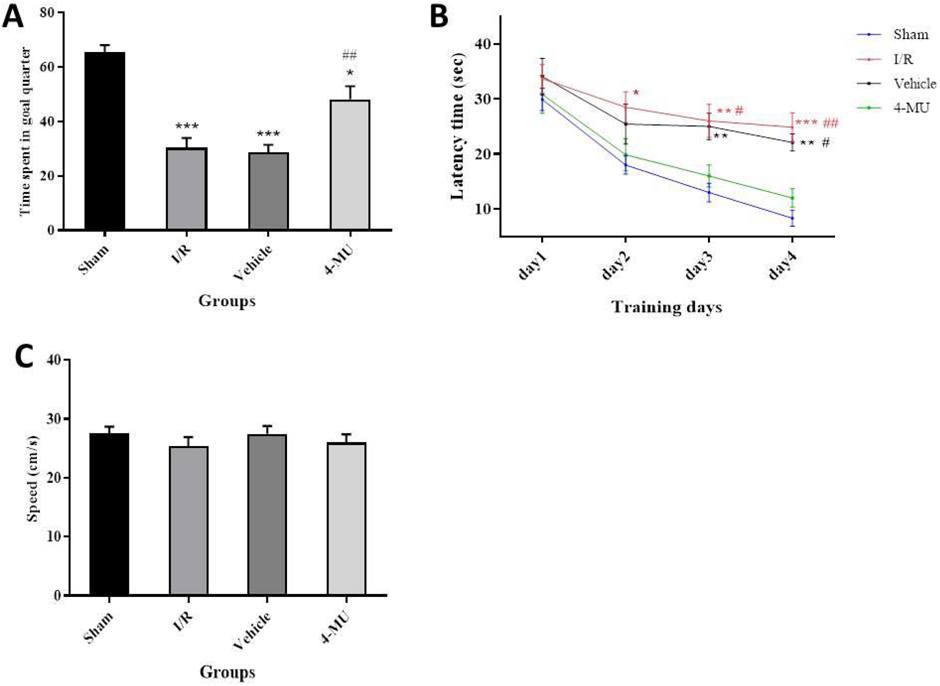
Impact of 4-MU on behavioral parameters in the Morris water maze (MWM) test (n□=□7 per group). **A)** The time spent in the goal quarter, **B)** Latency time, and **C)** Speed index. *□**□p□<□0.001, *□*□p□< □0.01, *□p□< □0.05 vs. sham; ## p□< □0.01 and #p□< □0.05 vs. I/R and vehicle cohorts). Data were expressed as mean ± □SEM.

### 3.4. Expression of HAS1 and HAS2

HAS1 and HAS2 as targets of 4-MU were found to be up-regulated following MCAO compared to the sham. Animals treated with 4-MU (25 mg/kg) exhibited decreased expression levels of both HAS1 and HAS2 in comparison with the I/R and vehicle cohorts (Figure 4 A-B).

**Figure 4.**
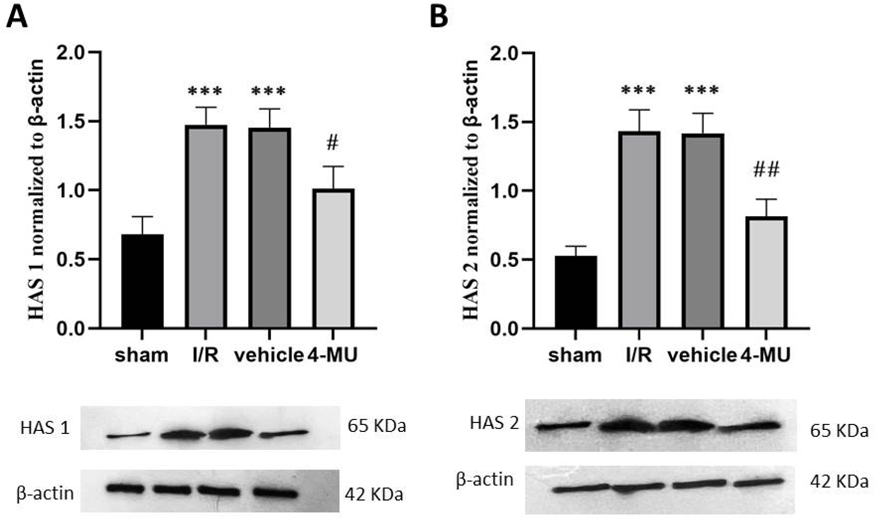
Western blot analysis for detection of the detection of the expression of A) HAS1, B) HAS 2 in brain tissues (n = 5). * * * p□<□0.001 vs. sham; ## p□<□0.01 and # p□< □0.05 vs. I/R and vehicle groups. Data were expressed as mean ± □SEM.

### 3.5. Neuronal density

Nissl staining of the CA1 region of the hippocampus revealed intact and tightly arranged neuronal cells with clear cytoplasm and nucleoli, exhibiting a monotonous staining pattern. In contrast, a scattered arrangement and nuclear deformation were observed in the I/R and vehicle cohorts compared to the sham. Treatment with 4-MU (25 mg/kg) significantly prevented cellular damage (Figure 5).

**Figure 5.**
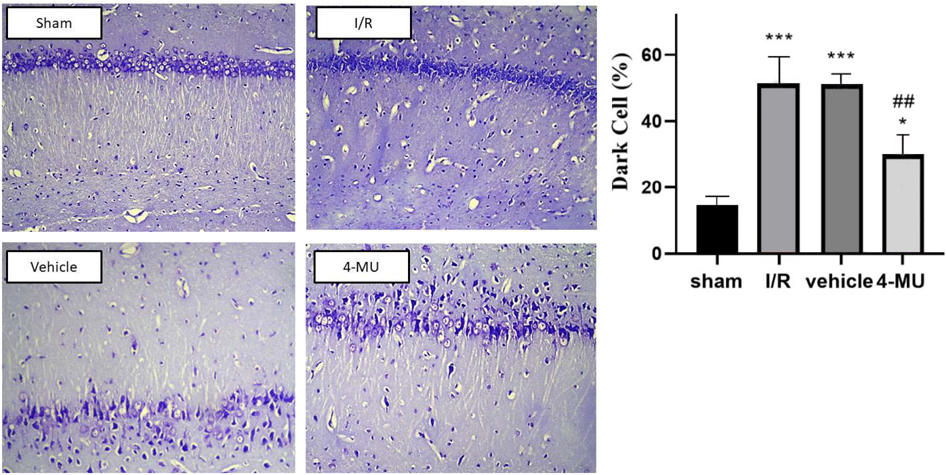
Representative images of Nissl staining in the CA1 region of the hippocampus with a scale bar of 50 μm) (n=5). ***p□<□0.001 vs. sham; ## p□<□0.01 and # p□< □0.05 vs. I/R and vehicle groups. Data were expressed as mean ± □SEM

## 4. Discussion

In addition to physical disability, cerebral I/R injury causes learning and memory impairments that heavily influence functional recovery post-stroke (6, 19). In the present study, our findings indicated that treatment with 4-MU (25 mg/kg) significantly attenuated infarct volume and reductions in spatial learning time (STL), along with elevated time in dark chamber, indicating learning and memory improvement. Consistent with our results, Dubisova et al. reported that oral administration of 4-MU attenuated perineuronal nets and improved recognition memory in mice (20).

Moreover, we found that beneficial effects of 4-MU in coping with I/R injury are associated with the inhibition of both HAS1 and HAS2. Emerging evidence demonstrates that 4-MU can reduce damage following I/R by suppressing oxidative stress and down-regulating TLR4/NF- κB/NLRP3 (21). Hyaluronic acid is a major ingredient of the brain extracellular matrix and a modulator of cell differentiation, migration, and angiogenesis. Prior investigations have found that hyaluronan accumulation in stroke-affected regions is linked to the overexpression of hyaluronidase-1 and 2 from one□hour to 21 days after stroke (22). A clinical study by Al’Qteishat et al. showed that the expression and concentrations of the enzymes responsible for HA synthesis and degradation markedly increased in post-mortem tissue and serum of patients at 1, 3, 7 and 14 days following stroke (6). Under homeostatic conditions, low HA-binding capacity has been reported for most immune cells. In inflammatory conditions, elevated HA binding of activated immune cells occurs via CD44 (20, 21). Increased expression of HAS isoenzymes (HAS1–3) during inflammation contributes to a predominance of low-molecular-weight HA, thereby activating macrophages through TLRs, which in turn results in elevated inflammation at the site of injury (22). These previous investigations are compatible with data herein indicating that tissue damage from MCAO resulted in up-regulation of HAS1 and HAS2, subsequently leading to obvious infarct volume and cell death in the CA1 region of the hippocampus. Such abnormalities were reversed by treatment with 4-MU (25 mg/kg), suggesting protective effects against cerebral I/R injury through inhibition of HAS1 and HAS2.

## 5. Conclusion

Ischemic stroke, a leading cause of morbidity and mortality worldwide, is characterized by the obstruction of blood flow to the brain, resulting in neuronal death and tissue damage. HAS enzymes are responsible for synthesizing HA, a glycosaminoglycan that plays a crucial role in regulating inflammation, cell migration, and tissue repair. 4-MU is a naturally occurring compound shown to inhibit HAS activity, thereby reducing HA production. The purpose of this study was to investigate the current understanding of the role of HAS in ischemic stroke and discuss the potential therapeutic effects of 4-MU on ischemic stroke through HAS inhibition. The neuroprotective effects of 4-MU in ischemic stroke are thought to be mediated through inhibiting HAS and subsequently reducing HA production. Additionally, 4-MU has been shown to reduce infarct size, improve neurological function, and enhance cognitive recovery in animal models of MCAO. In this study, we found that 4-MU may be a promising candidate for reducing cerebral I/R injury and improving learning and memory impairments in a rat model of ischemic stroke. It seems likely that the neuroprotective impact of 4-MU in coping with cerebral I/R injury is related to suppressing the activities of HAS1 and HAS2.

## Supporting information

author's affiliations and Declarations

## Abbreviations

4-MU: 4-methylumbelliferone
MCAO: Middle Cerebral Artery Occlusion
HAS: Hyaluronan Synthase
I/R: Ischemia-reperfusion

---

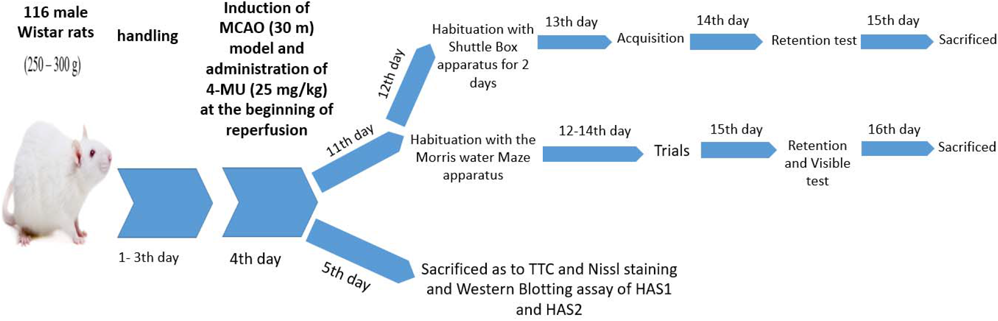

