## Supplementary material for "4-Methyllumifrone (4-MU) can improve learning and memory after cerebral ischemia/reperfusion injury in rats": author's affiliations and Declarations

Author’s affiliation:

1. Physiology Research Center, Iran University of Medical Sciences, Tehran, Iran.
2. Physiology Research Center, Iran University of Medical Sciences, Tehran, Iran.
3. Department of Physiology, School of Medicine, Iran University of Medical Sciences, Tehran, Iran
4. Laser Research Centre, Faculty of Health Science, University of Johannesburg, Doornfontein 2028, South Africa.
5. Physiology Research Center, Iran University of Medical Sciences, Tehran, Iran.
6. Physiology Research Center, Iran University of Medical Sciences, Tehran, Iran.

**Corresponding authors:**

Nahid Aboutaleb,

**Declarations**

**Ethics approval and consent to participate**

This proposal was approved by IRAN University of Medical Sciences Ethics Approval Center with code number IR.IUMS.REC.1401.378.

**Consent for publication**

Not applicable.

**Clinical trial number**

Not applicable.

**Availability of data and material**

The findings of this study are derived from data that are available upon request from the corresponding authors.

**Competing interests**

The authors declare no conflicts of interest.

**Funding**
FR was supported by the IRAN University of Medical Sciences, Grant no. 1401.378.

**Authors' contributions**

N.A and F.R participated in the design and interpretation of the study, data analysis, and the review of the manuscript. SH.T, H.M-J , E.K and F.R conducted the experiment, collected the tissue samples, and were responsible for the data analysis. SH.T performed histopathological analyses. SH.T and F.R wrote the manuscript, and all authors reviewed, read, and approved the article.

**Acknowledgements**
Not Applicable.
